# Focused ultrasound-enhanced carboplatin delivery improves radiotherapy response and survival in a preclinical glioblastoma model

**DOI:** 10.64898/2026.11.17.766937

**Authors:** Sébastien Rigollet, Sarvenaz Keshmiri, Sarra Dbira, Marie D’Allemagne, Thomas Ador, Alexandre Leboucher, Vasile Stupar, Benjamin Lemasson, Anthony Delalande, Jean-François Adam, Anne-Laure Bulin, Benoît Chovelon, Erik Dumont, Jean-Luc Ravanat, Emmanuel L. Barbier, Hélène Elleaume

## Abstract

**Rationale:** Glioblastoma is highly resistant to conventional therapies due to the limited penetration of chemotherapeutics across the blood-brain and blood-tumor barriers (BBB, BTB). Although platinum-based drugs show strong cytotoxic and radiosensitizing properties in several cancers, their inability to achieve therapeutic concentrations within the central nervous system (CNS) severely limits their efficacy in brain tumors. This study investigated whether focused ultrasound (FUS)-mediated BBB/BTB disruption combined with intravenous carboplatin and stereotactic radiotherapy could improve drug delivery, tumor control, and survival in a preclinical glioblastoma model.

**Methods:** F98 glioma-bearing rats were treated with a single dose of 15 Gy radiation therapy (RT), carboplatin only, carboplatin + RT, FUS + carboplatin or FUS + carboplatin + RT. FUS-mediated permeabilization was confirmed by magnetic resonance imaging (MRI), and platinum accumulation was quantified by inductively coupled plasma mass spectrometry and synchrotron X-ray fluorescence. Tumor progression and microenvironmental changes were monitored using multiparametric MRI (diffusion, perfusion, and vascular permeability), followed by voxel-wise clustering analysis and histopathological validation.

**Results:** The FUS + carboplatin + RT group demonstrated the strongest therapeutic benefit, compared to all other treatment groups, yielding up to a 122% increase in median survival relative to the untreated group. This treatment also reduced necrotic core expansion and induced a transient increase in tumor perfusion, suggesting improved vascular function. FUS enhanced carboplatin delivery to both the tumor and peritumoral brain, with platinum distribution closely correlating with MRI signal enhancement. Voxel-wise clustering of multiparametric MRI identified biologically distinct tumor regions that were subsequently validated by immunohistochemistry.

**Conclusion:** The combination of FUS-mediated carboplatin delivery and radiotherapy represents a safe, non-invasive strategy that enhances drug delivery and improves survival in an aggressive preclinical model of glioblastoma. By overcoming a major limitation of current therapies, this approach shows promising potential for clinical translation.

## Introduction

Glioblastoma represents the most prevalent primary brain tumor in adults [1,2], characterized by aggressive infiltration, rapid progression and marked resistance to therapy [3]. Its incidence has steadily increased over the past decade, from 2.1 to 3 cases per 100,000 individuals [4], highlighting the urgent need for more effective therapeutic strategies. The current standard of care, established by Stupp et al. in 2005, consists of maximal safe surgical resection followed by radiotherapy (60 Gy delivered in 30 fractions) with concomitant daily temozolomide chemotherapy, followed by six cycles of adjuvant temozolomide chemotherapy. This regimen achieved a median overall survival of 14.6 months, compared with 12.1 months for radiotherapy alone, in a landmark phase III clinical trial [5]. Despite subsequent advances in neurosurgical techniques and the introduction of tumor-treating fields (TTFs), median overall survival remains approximately 20 months after diagnosis [6].

Radiotherapy is a cornerstone of glioblastoma treatment, targeting both the surgical cavity and the peritumoral zone, which harbors infiltrative tumor cells extending beyond the visible tumor core. Dose escalation in these functionally critical regions is precluded by the risk of radionecrosis. As a result, 80-90% of recurrences occur within the peritumoral zone [7], and the current standard of care is therefore not curative.

Improving radiotherapy efficacy through radiosensitizing agents is therefore a central research objective. The therapeutic benefit of most chemotherapeutic agents is, however, substantially limited by the heterogeneous permeability of the blood–tumor barrier (BTB) within the tumor core and, more critically, by the intact blood–brain barrier (BBB) in the peritumoral zone, precisely where recurrences predominantly arise [9]. Temozolomide remains the standard chemotherapy agent, not because of its superior cytotoxic potency, but because of its favorable physicochemical properties that enable efficient penetration across the BBB [8,9]. By contrast, platinum-based compounds offer greater intrinsic antitumor efficacy and potent radiosensitizing properties (augmenting RT-induced DNA double-strand breaks, inhibiting DNA repair, and triggering oxidative stress and immunogenic cell death [10]), yet their clinical utility in central nervous system malignancies is severely limited by poor BBB penetration [10].

To bypass the BBB, convection-enhanced delivery (CED) of platinum agents has been explored. Preclinical studies using CED of carboplatin or cisplatin, in combination with radiotherapy have demonstrated complete tumor regression and long-term survival in F98 glioma models [11]. However, the clinical translation of CED remains limited by its highly invasive nature and technical challenges including catheter misplacement and heterogeneous drug distribution [12].

More recently, focused ultrasound (FUS) combined with microbubbles has emerged as a promising non-invasive strategy for transient BBB/BTB disruption and enhanced drug delivery. Clinical trials have demonstrated its feasibility in glioblastoma patients (NCT05902169, NCT06329570, NCT04988750) [13–18], and recent data support the benefit of large-volume peritumoral BBB opening: the Sonocloud-9 device combined with carboplatin showed encouraging results in recurrent glioblastoma [14], large-volume BBB disruption enabling the delivery of either carboplatin or temozolomide was demonstrated to be safe in glioblastoma patients [19], and a phase 1/2 trial with transcranial FUS and temozolomide reported improved outcomes in high-grade glioma [20]. Large-volume sonications have been employed safely across multiple preclinical contexts, including whole-brain BBB opening for AAV gene delivery that was associated with survival improvements [21] and repeated large-volume openings in healthy rats without significant structural damage or glial reactivity [22]. Importantly, when large volumes are achieved through multiple overlapping foci, the BBB closure time and opening dynamics remain independent of the sonicated volume [23]. In preclinical models, however, FUS-enhanced temozolomide delivery has yielded inconsistent survival benefits [24,25], possibly because disrupting the BBB at the tumor core (where the barrier is already compromised) confers limited additional advantage. By contrast, FUS-mediated delivery into the intact peritumoral BBB, the very zone where recurrences arise in over 80% of cases, represents a more mechanistically sound target. Carboplatin is particularly well-suited to exploit this opportunity: beyond its antitumor potency, it is a potent multi-modal radiosensitizer, augmenting RT-induced DNA double-strand breaks, inhibiting DNA repair pathways, and triggering oxidative stress and immunogenic cell death [10]. Delivering carboplatin into the peritumoral brain with FUS followed by radiotherapy would therefore simultaneously address two critical failure modes of current treatment (i.e., insufficient drug penetration and inadequate radiosensitization) in the zone where the disease recurs. Despite this compelling rationale, this triple combination has not yet been evaluated, to our knowledge.

Recent advances in multiparametric magnetic resonance imaging (MRI) have enabled non-invasive monitoring of key tumor features, including cellularity (with the apparent diffusion coefficient (ADC)), cerebral blood flow (CBF), and vascular permeability (with dynamic contrast-enhanced MRI (DCE-MRI)) [26]. These biomarkers provide complementary information for characterizing therapeutic effects in glioblastoma across preclinical and clinical settings [26,27]. Multiparametric clustering approaches further integrate these parameters into a spatially resolved overview of tumor heterogeneity and have been shown to stratify non-enhancing T_2_-FLAIR hyperintense regions into prognostic subtypes correlated with progression-free survival in glioblastoma patients and to identify distinct tumor habitats [28–31].

Here, we evaluate the therapeutic efficacy of combining intravenous (i.v.) carboplatin delivery with FUS-mediated BBB/BTB disruption and stereotactic radiotherapy in the syngeneic immunocompetent F98 rat glioma model, a highly relevant preclinical system owing to its low immunogenicity, highly infiltrative growth pattern, and morphological similarity to human glioblastoma [32–35]. This strategy is based on three interconnected rationales: (i) delivering chemotherapy to the peritumoral zone, where the blood-brain barrier (BBB) remains intact and where recurrences predominantly arise; (ii) exploiting the radiosensitizing properties of carboplatin to enhance the efficacy of radiotherapy; (iii) leveraging the non-invasive, clinically translatable nature of focused ultrasound (FUS). We first quantified platinum biodistribution using inductively coupled plasma mass spectrometry (ICP-MS) and synchrotron X-ray fluorescence (SR-XRF) imaging to assess the impact of FUS-mediated BBB/BTB opening on carboplatin delivery. Tumor progression and survival were then assessed across treatment groups. Longitudinal multiparametric MRI tracking water diffusion, perfusion and BBB permeability provided non-invasive characterization of the tumor microenvironment, and voxel-wise clustering was applied to identify biologically distinct tumor subregions across groups.

## Materials and Methods

### Animal ethics

The experiments were performed using 99 male Fischer rats (Charles River laboratory, Lyon, France), aged 7 to 8 weeks at the time of arrival. All animal protocols were approved by the Ethics Committee and the French Ministry of Research and Health (authorization APAFIS #48225) and conducted in accordance with institutional and national guidelines for animal care. Animal weight and general health status were monitored daily. During all the experimental procedures, the animals were anesthetized with isoflurane (2 – 3%) in air.

### Study design overview

The study comprised three experimental sub-studies: a therapeutic efficacy study (47 animals, six treatment groups, longitudinal multiparametric MRI follow-up), an acute safety study (18 animals, assessed at 48 h post-sonication), and a platinum biodistribution study (22 animals, assessed by ICP-MS and synchrotron X-ray fluorescence at 1 h and 24 h post-injection). Animal numbers specific to each sub-study are provided in the corresponding sections below and recapitulated in Table S1.

### Safety study

Eighteen animals were included to evaluate the acute safety profile of FUS-mediated BBB/BTB opening. MRI was performed 48 h after sonication to assess BBB closure, followed by brain harvesting for histological analysis. Brain sections were stained with hematoxylin and eosin (H&E) and examined for evidence of tissue damage, including hemorrhage. This cohort also completed the MRI dataset: multiparametric MRI was acquired at 2 weeks post-implantation in the untreated group (N = 15), providing additional tumor MRI data at this timepoint e.g. the time of treatment.

### Platinum biodistribution study

Twenty-two animals were included in the biodistribution study, which comprised two complementary approaches: (A) ICP-MS for quantitative assessment of platinum concentration in brain tissue samples, and (B) SR-XRF imaging for two-dimensional mapping of platinum spatial distribution in brain sections.

- A) ICP-MS quantification: Animals were assigned to the following groups: untreated (N = 2); carboplatin, 1 h after injection (N = 3); carboplatin, 24 h after injection (N = 3); FUS + carboplatin, 1 h after treatment (N = 3); FUS + carboplatin, 24 h after treatment (N = 3).
- B) X-ray fluorescence imaging (1 h after treatment): Animals were assigned to the following groups: untreated (N = 2); carboplatin only (N = 3); FUS + carboplatin (N = 3).

At the predefined time points after treatment, blood samples were collected immediately before euthanasia. The brain and peripheral organs, including the liver, kidneys, heart, and spleen, were harvested post-mortem, briefly immersed in pre-cooled isopentane (–80⍰°C) to ensure rapid freezing, and subsequently stored at –80⍰°C until further processing.

### Therapeutic study

Forty-seven animals were included in the therapeutic study. Two weeks following tumor implantation, animals were anesthetized and underwent MRI to confirm tumor growth. Based on these initial MRI data, animals were randomly assigned to treatment groups to ensure comparable tumor burden across conditions: untreated (N = 11), carboplatin only (N = 5), radiotherapy (RT) only (N = 6), carboplatin + RT (N = 5), FUS + carboplatin (N = 10), and FUS + carboplatin + RT (N = 10). Animals in the FUS + carboplatin and FUS + carboplatin + RT groups received i.v. microbubbles followed by FUS-mediated BBB/BTB opening, as described in the corresponding section, immediately followed by carboplatin administration.

Animals in the FUS + carboplatin + RT group received radiotherapy 24 h after the FUS-mediated BBB/BTB opening and carboplatin administration. All animals underwent MRI examinations to monitor tumor evolution (Figure 1.A). Body weight was recorded daily, and any signs of adverse effects affecting general appearance or behavior were documented. Euthanasia was performed when animals exhibited markedly decreased activity or body weight loss exceeding 20% over one week. Brains were collected immediately after euthanasia, briefly immersed in pre-cooled isopentane (– 80⍰°C) to ensure rapid freezing, and then stored at –80⍰°C until further processing for histological analysis.

**Figure 1.**
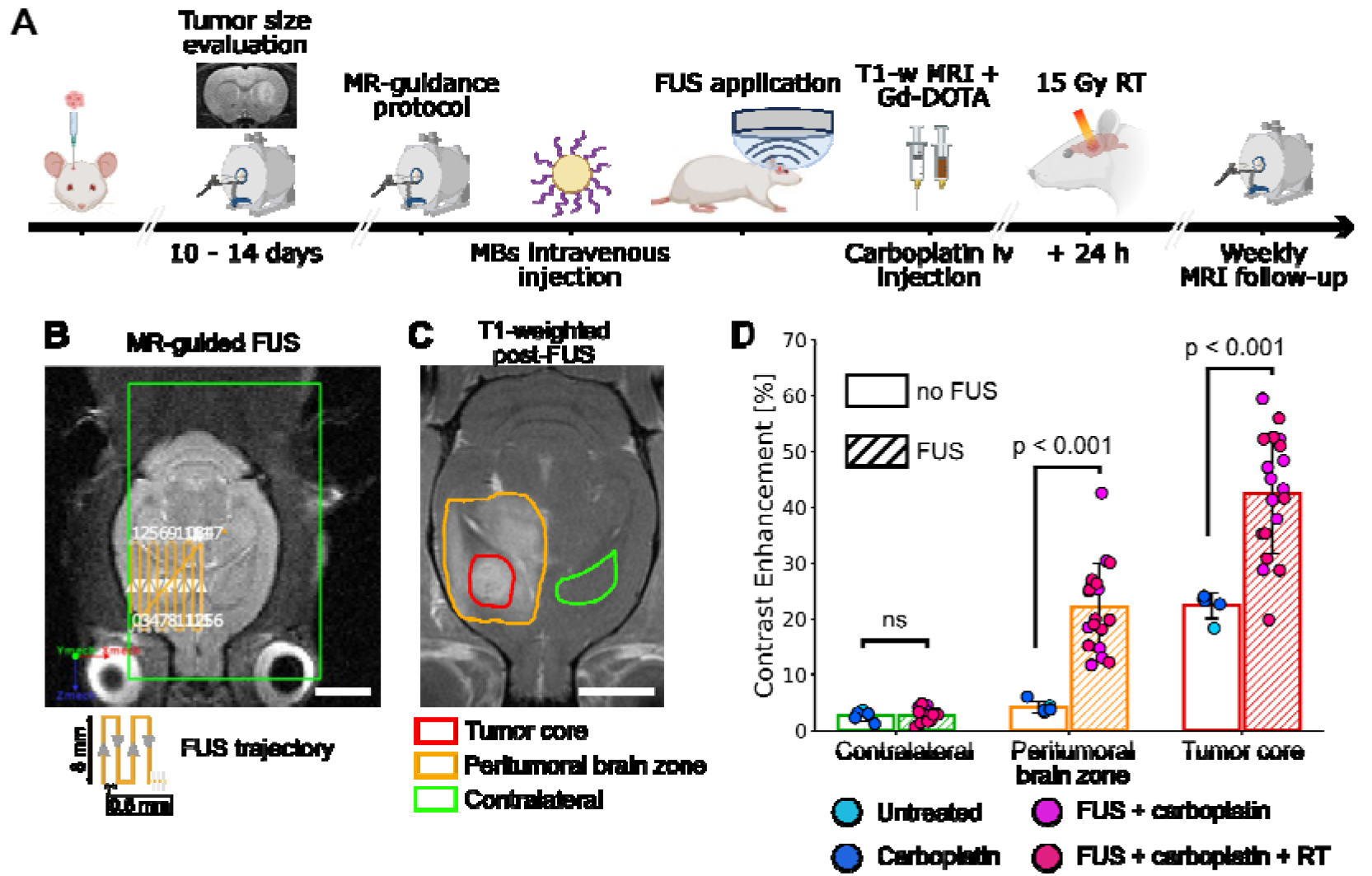
BBB/BTB opening using MR-guided FUS and microbubbles. A) Experimental timeline: following intracerebral injection of 10^3^ 10 F98 cells, tumor size was evaluated before any procedure. The BBB/BTB was permeabilized using MR-guided FUS in combination with microbubbles. Following the confirmation of FUS-induced BBB/BTB opening by contrast-enhanced MRI, carboplatin was injected intravenously. A radiation dose of 15 Gy was delivered 24 h after FUS. B) Screenshot of Thermoguide® software showing the T_2_-weighted anatomical image with the overlaid 8 × 8 mm² raster scan zone (yellow) used to target the tumor (scale bar, 5 mm). C) Contrast-enhanced T_1_-weighted MR image acquired after FUS and Gd-DOTA injection. Three anatomical regions of interest (ROIs) were delineated: contralateral striatum (green), peritumoral brain zone (orange), and tumor core (red) (scale bar, 5 mm). D) Gd-DOTA signal enhancement (%) in each MRI-derived ROI for non-FUS (cyan, blue) and FUS (hatched bars, purple, pink) animals. P-values were derived from Mann-Whitney U tests (ns: not significant). Data are presented as mean ± SD.

### F98 rat model

F98 rat glioma cells, obtained from ATCC, were cultured in T75 flasks in high-glucose DMEM (GlutaMAX, Thermo Fisher Scientific) supplemented with 10% fetal bovine serum and 1% penicillin/streptomycin under standard conditions (37⍰°C, 5% CO₂). Upon reaching 80–90% confluence, cells were trypsinized, washed, and resuspended in serum-free DMEM. Cell concentration was determined using a Malassez hemocytometer and adjusted to 2.5⍰×⍰10⁵ cells/mL. Glioma cells were orthotopically inoculated into the brain of male Fischer F344 rats (Charles River®, France) following the procedure described previously [36]. The rats were anesthetized with isoflurane and immobilized in a stereotactic frame (David Kopf Model 902). After shaving, the skull was exposed through a scalp incision, and a small hole was made using a 23 G needle, at 0Tmm (anteroposterior) and 3.5⍰mm (lateral), relative to the bregma in the right frontal hemisphere. A 32 G Hamilton needle mounted on a 25 µL Hamilton sealed syringe was stereotactically inserted into the injection site at a depth of 5.5 mm from the skull surface. A 4⍰µL volume of F98 cell suspension (containing 1000 cells) was delivered at a rate of 1⍰µL/min using a syringe pump (KDS310; Geneq, Inc., Montréal, Quebec, Canada) to minimize backflow. Following tumor cell implantation, the needle was left in place for 1 min before being slowly withdrawn over approximately 1 min. The skull hole was then closed with bone wax, and the skin was sutured.

### MR-guided focused ultrasound procedure

Anatomical T_2_-weighted images were acquired to confirm tumor presence, dimensions and localization. FUS was applied using an MR-guided device (Image Guided Therapy, Pessac, France) equipped with a 1.5 MHz spherically focused transducer (Imasonic, Voray-sur-l’Ognon, France). The transducer consists of seven concentric elements allowing dynamic focusing at varying depths. The acoustic beam profile of the transducer was previously characterized and calibrated [37], showing a “rice grain”-shaped focal zone with dimensions of 1.15 × 1.15 × 5.5 mm^3^ (full width at half maximum). Prior to the sonication procedure, animals were anesthetized, and the scalp was shaved and depilated to minimize ultrasound attenuation. Ultrasound gel was applied to the scalp to ensure efficient acoustic coupling between the transducer’s membrane and the brain. The cradle was then placed in the 4.7T MRI scanner (see MRI acquisition section below for details) and T_1_-weighted axial and coronal images were acquired to localize the transducer relative to the rat brain. The images were imported into Thermoguide® software where an 8 × 8 mm² raster scan was defined, centered on the tumor identified as hyperintense on T_2_-weighted images; see Figure 1.B. The scan encompassed both the tumor core and the peritumoral brain region to enable BBB and BTB permeabilization. Before the sonication protocol was initiated, 1.5 × 10^8^ lipidic microbubbles diluted in 100 μL of HEPES buffer (10 mM, pH = 7.4) were i.v. injected via the tail vein and flushed with 50 μL of saline (0.9% NaCl, Versol®). Details on microbubbles composition, activation and characteristics can be found in a previous study [38]. Sonication was performed using continuous wave ultrasound at a peak negative pressure of 0.44 MPa estimated in situ [39]. This pressure was selected based on previous findings demonstrating effective carboplatin delivery and improved survival in the F98 rat glioma model [40]. The transducer moved at a velocity of 10 mm/s, with a 50 ms pause between directional changes to prevent ultrasonic energy accumulation. The total duration of sonication per animal was approximately 90 s.

### Carboplatin administration

Carboplatin (Teva Pharmaceuticals; 10 mg/mL, 50 mg/kg) was administered intravenously via the tail vein over approximately 90 s, following confirmation of FUS-induced BBB/BTB opening (i.e., within 30 min after FUS application). This dose corresponds to a previously reported human equivalent dose of 300 mg/m^2^ [40].

### Radiotherapy protocol

Radiotherapy was performed using a Small Animal Radiation Research Platform (SARRP; Xstrahl Ltd., UK) at the SAXO-IRMaGe facility in Grenoble, France. Rats were anesthetized and positioned on a dedicated cradle within the SARRP system. A cone-beam computed tomography (CBCT) scan was first acquired using the onboard imager of the SARRP. The T_2_-weighted MRI images were co-registered to the CBCT images using the Muriplan® treatment planning system. Dose planning calculations were performed to deliver 15 Gy to the tumor center, while ensuring that more than 70% of the tumor volume and a 3 mm safety margin around it received a dose exceeding 14.5 Gy. Radiation was delivered through a 10 × 10 mm² collimator to guarantee precise dose administration across the entire tumor volume (see Figure S1). Treatment was administered using an arc ranging from −75 to + 75°, with the following parameters: tube voltage of 220 kV, tube current of 13 mA, source-to-surface distance of 35 cm, a 0.15 mm Cu filter, and a dose rate of 3 Gy/min. The irradiation time was approximately 350 s per animal. In the FUS + carboplatin + RT and carboplatin + RT groups, radiotherapy was delivered 24 h after intravenous carboplatin administration. In the RT-only group, irradiation was performed the day after enrollment in the therapeutic protocol, based on tumor volume as previously described.

### Magnetic Resonance Imaging

MRI was performed at 4.7 T (Avance III console, Bruker; IRMaGe facility, Grenoble, France) using a volume coil for transmission (transmit coil, Bruker) and a surface receiver coil (Bruker) under 1.5-2% isoflurane anesthesia.

#### Tumor assessment

Tumor detection and initial volumetric evaluation were performed 14 days after tumor implantation using anatomical MR T_2_-weighted imaging. Tumor segmentation was manually performed on T2-weighted images, and tumor volumes were calculated by summing tumor areas across consecutive slices multiplied by slice thickness.

#### MRI follow-up

A multiparametric MRI examination was performed two weeks after treatment and weekly thereafter to monitor tumor progression and assess therapeutic response. The imaging sessions included the acquisition of the following sequences: high-resolution T_2_-weighted TurboRARE, inversion recovery T_1_ mapping, pseudo-continuous arterial spin labeling (pCASL), diffusion-weighted imaging (DWI), dynamic contrast-enhanced MRI (DCE-MRI), and an additional inversion recovery slice for sequence quality control. Gd-DOTA contrast agent (Dotarem®, Guerbet, Villepinte, France) was i.v. injected at a dose of 200 µmol/kg, one minute after the initiation of the DCE sequence. This imaging protocol was repeated weekly for all animals. Detailed MRI acquisition parameters are summarized in Table 1.

**Table 1.** Summary of the MRI sequences. The multiparametric protocol included anatomical, perfusion, diffusion, and contrast-enhanced sequences to characterize tumor morphology and therapeutic response. Abbreviations: FOV, field of view; Inv. Eff., pCASL inversion efficiency; pCASL, pseudo-continuous arterial spin labeling; T_1_-MTI, T_1_ mapping with multiple inversion times; LD, labeling duration; PLD, post-labeling delay; T_acq_, acquisition time; TE, echo time; TI, inversion time; TR, repetition time.

| **Sequence** | **TR/TE [ms]** | **FOV [mm<sup>2</sup>]** | **matrix size** | **voxel size [µm<sup>3</sup>]** | **Other parameters** | **T<sub>acq</sub>** |
|---|---|---|---|---|---|---|
| T<sub>2</sub> TurboRARE | 2200/36 | 30 × 30 | 256 × 256 | 117 × 117 × 1000 | RARE-factor = 8 | 3 min 31 s |
| DTI-EPI | 2200/33 | 30 × 30 | 128 × 128 | 234 × 234 × 1000 | b-values = 0, 800 s.mm<sup>-2</sup> | 3 min 31 s |
| Inv. Eff. | 225/3.6 | 30 × 30 | 256 × 256 | 117 × 117 × 1000 | 1 mm-thick, single slice | 3 min 30 s |
| T<sub>1</sub>-MTI | 10000/19 | 30 × 30 | 128 × 128 | 234 × 234 × 1000 | 18 TIs = {30; 10000 ms} | 4 min |
| pCASL | 4000/21 | 30 × 30 | 128 × 128 | 234 × 234 × 1000 | 30 repet., LD = 3000 ms, PLD = 300 ms | 4 min |
| DCE-MRI | 800/4 | 30 × 30 | 128 × 128 | 234 × 234 × 1000 | Gd-DOTA<br>injection at 1 min | 6 min 20 s |

#### MRI Data processing

ADC maps were calculated using ParaVision 6.0.1. The CBF parameter was computed using the following three sequences: the inversion efficiency slice, the T_1_ map and the pCASL dataset. Data processing was performed according to previously published data [37,41]. MRI data were analyzed and processed using MP3 software [42]. Images were registered using FSL FLIRT [43] to the first-week acquisition as a within-animal reference for longitudinal analysis. The area under the curve (AUC) over the duration of the imaging time and the time-to-peak (TTP) in seconds were calculated from the DCE-MRI signal enhancement curves.

#### Quantitative image analysis

In the therapeutic cohort, full multiparametric MRI was not acquired at the treatment timepoint (week 2 post-implantation) to avoid combining the imaging session with the treatment procedures on the same day. To establish the k-means clustering framework at this timepoint, baseline multiparametric MRI data from the untreated control subgroup of the safety cohort (N = 15; see Safety study and Table S1) were used to obtain multiparametric MRI data at 2 weeks post-implantation.

First, a region-wise analysis was performed using manually delineated regions on the reconstructed MRI contrasts. A region of interest (ROI) was initially drawn on T2-weighted images to estimate the total tumor volume, defined by the hyperintense signal encompassing the tumor core, the peritumoral brain zone, and the surrounding edema. Unsupervised clustering was performed on the T_2_-weighted-based ROI using the k-means method [44]. The analysis incorporated ADC, CBF, T_1_, AUC, and TTP parameters and was conducted voxel-wise across the entire tumor volume defined on T2-weighted images. Non-physiological voxels were excluded using the following thresholds: 0 < ADC < 2500 µm²/s and 0 < CBF < 300 mL/min/100 g. For the other parameters, no thresholds were applied, as non-physiological values were not identified. Voxels meeting any exclusion criterion were removed from all maps. The remaining data were then standardized using the mean and standard deviation across all voxels to minimize scale differences between parameters. A total of 77 MRI time points, corresponding to the different imaging sessions across all animals and experimental timepoints, were included in the analysis. Voxels from all animals were pooled and analyzed using the k-means algorithm from the scikit-learn library in Python (version 3.10.0). A total of 1,105,471 voxels were included. The optimal number of clusters was determined using the elbow method in combination with the silhouette score, resulting in five clusters [45,46].

### Platinum biodistribution methods

#### Inductively Coupled Plasma Mass Spectrometry

The brain was dissected into two distinct regions—the contralateral hemisphere and the tumor-bearing hemisphere. Peripheral organs were sampled by collecting approximately 200 to 300 mg of tissue per specimen. Samples were mineralized under atmospheric pressure in nitric acid for 24 h at room temperature, followed by three successive 8 h phases in an oven at 50°C, over three consecutive days. The resulting mineralized samples were diluted to a final nitric acid concentration of 1% prior to analysis. Platinum calibration standards were prepared in 1% (v/v) nitric acid. Platinum concentrations were determined using quadrupole ICP-MS (Perkin Elmer NexION 2000, Waltham, MA, USA). The ^195^Pt was quantified and ^103^Rh was used as an internal standard.

#### Synchrotron Radiation X-ray Fluorescence Imaging

Synchrotron-based X-ray fluorescence (SR-XRF) enables highly sensitive and spatially resolved elemental mapping, with micrometer scale resolution. It allows the simultaneous detection and quantification of multiple trace elements in biological tissues without the need for chemical labeling, making it particularly well-suited for investigating the biodistribution of metal-based drugs. In the present study, SR-XRF was used to quantify and visualize the spatial distribution of platinum in the brains of F98 glioma-bearing rats following treatment with carboplatin, administrated either intravenously alone or in combination with FUS and microbubbles to enhance drug delivery across the BBB and BTB. To assess early platinum distribution, animals were euthanized 1 h after carboplatin administration. Organs were rapidly collected, snap-frozen, and sectioned at 40Tµm thickness using a cryotome maintained at –20T°C. Sections were collected at multiple rostro caudal levels of the brain, as well as at representative orientations of peripheral organs, including the kidney, spleen, and liver. All tissue sections were mounted on RNAse-free membrane slides with aluminum frames (MMI Membrane Slides, ref. 50102, Molecular Machines & Industries, Germany) and stored at −20T°C until analysis.

SR-XRF imaging was performed at the ID15A beamline of the European Synchrotron Radiation Facility (ESRF, Grenoble, France) under proposal number Elleaume-MD1425. The X-ray beam energy was set to 100 keV (350 eV bandwidth) and focused to a spot size of 18⍰×⍰30⍰μm² (vertical × horizontal), a photon flux of approximately 2 x 10^11^ photons/s at the focal spot. Fluorescence signals were recorded using a single thermoelectrically cooled Canberra (Mirion Technologies) GeCMOS2 detector positioned at 90° relative to the incident X-ray beam (active diameter: 10 mm; crystal thickness: 6 mm) and equipped with a custom-built tungsten-lead collimator designed to minimize background noise and enhance detection sensitivity. The sample-to-detector distance was 6.5 cm. The sample slide was mounted vertically at an angle of 7° relative to the detector axis to optimize fluorescence collection. The experimental setup is illustrated in Figure S2.A. Additional technical details regarding the ID15A beamline setup have been previously described [47]. Samples were raster-scanned with an effective pixel size of 200 × 200 μm², with an acquisition time of 10 s per pixel. Imaging of a single brain section typically required approximately 8 h of acquisition time.

SR-XRF image Analysis: All SR-XRF data processing and spectral fitting were performed using PyMCA, version 5.9.4 [48]. To quantify platinum levels in biological samples, a multi-step calibration and modelling approach was applied:

1. **Primary calibration:** Calibration was performed using fluorescence maps acquired from the certified reference sample RF11-200-s3235-09 (AXO-Dresden, Germany), containing known concentrations of multiple elements (Pb, La, Pd, Mo, Cu, Fe, Si). The incident beam intensity and experimental geometry were calibrated by modelling the measurement setup and simulating fluorescence spectra based on this reference.
2. **Secondary calibration:** Additional calibration was performed using spectra acquired from glass capillaries (2 mm internal diameter) filled with known carboplatin concentrations (10, 100, and 1000 µg/g), corresponding to platinum concentrations of 5.25, 52.55, and 525.50 µg/g, respectively. These measurements were used to verify linearity and improve quantitative accuracy (Figure S2.B).
3. **Pixel-wise spectral modelling:** Using the calibration parameters obtained from the AXO reference material, theoretical spectra were simulated for each pixel of the biological samples to estimate local elemental concentrations (Figure S2.C).
4. **Conversion to volumetric concentration:** Mass per surface area (g/cm²) values obtained from SR-XRF analysis were converted into volumetric concentrations (ppm) using a section thickness of 40⍰µm and assuming a tissue density of 1⍰g/cm³.

This approach enabled the generation of quantitative platinum distribution maps throughout tissue sections. Images were then analyzed with ImageJ, and ROIs were drawn in the contralateral and tumor-bearing hemisphere to perform further analysis.

### Histology

For histological analysis, brains were harvested, snap frozen in isopentane, and stored at −80°C. Cryosections (10 µm thick) spanning different levels along the anteroposterior axis were obtained using a cryotome settled at −20°C. To enable multiple analyses, sections dedicated to different staining procedures, including hematoxylin and eosin (H&E) and immunohistochemical labeling of vascular and proliferation markers, were collected in an interleaved manner. For immunohistochemistry, sections were fixed in paraformaldehyde (Fluka Chemie AG, Buchs, Switzerland) and incubated overnight at 4°C with the following primary antibodies: goat anti-collagen IV (ColIV) (1:1000, AO305, Southern Biotech) to assess the basal lamina; mouse anti-CD31 (1:1000, MAB1393, Merck Millipore) to label endothelial cells; and rabbit anti-Ki67 (1:250, MA5-14520, ThermoFisher) to evaluate cell proliferation. After primary antibody incubation, sections were incubated with species-specific fluorophore-conjugated secondary antibodies: donkey anti-goat A546 (1:1000, Invitrogen, A11056), donkey anti-mouse A488 (1:1000, A21202, Invitrogen) and goat anti-rabbit A568 (1:200, A11036, Invitrogen). Sections were then counterstained with DAPI (4T,6-diamidino-2-phenylindole, ThermoFisher) for nuclear visualization, mounted with coverslips, and rinsed in PBS containing 0.1% Tween-20. Images were digitized using the Axioscan scanner (Zeiss, Oberkochen, Germany) at the PIC-GIN photonic imaging platform (Grenoble, France).

### Statistical analysis

Survival data were analyzed using the Kaplan-Meier method and compared between groups using the log-rank test. For MRI-derived parameters (tumor ADC, CBF, and their volumetric subregions), between-group differences at each timepoint were assessed using a one-way ANOVA. When a significant global effect was detected (*p* < 0.05), pairwise post-hoc comparisons were performed using the Mann-Whitney U test. For platinum biodistribution data, group differences were evaluated using two-sided paired t-tests. The temporal evolution of k-means cluster frequencies across treatment groups was assessed by one-way ANOVA at each timepoint. All statistical analyses were performed in Python 3.10 (scipy and statsmodels libraries). Data are presented as mean ± SD unless otherwise stated. A *p*-value < 0.05 was considered statistically significant.

## Results

### MR-guided FUS induces localized and safe BBB/BTB opening

In the FUS + carboplatin and FUS + carboplatin + RT groups, transient BBB/BTB opening was induced using MR-guided FUS in combination with i.v. administered microbubbles (Figure 1.A–B). Immediately after sonication, Gd-DOTA was injected intravenously and contrast-enhanced T₁-weighted MRI was performed to assess BBB/BTB permeabilization. The permeability was measured in three regions of interest: the contralateral striatum, the peritumoral brain zone, and the tumor core (Figure 1.C).

FUS-treated animals showed significantly higher Gd-DOTA enhancement than non-FUS controls in both the tumor core (FUS-treated: 42.5 ± 10.7% vs. non-FUS: 22.4 ± 2.3%; *p* < 0.001, Mann-Whitney U) and the peritumoral brain zone (FUS-treated: 22.2 ± 7.7%; non-FUS: 4.2 ± 1.1%; *p* < 0.001, Mann-Whitney U), while enhancement in the contralateral striatum remained below 3% across all groups (ns), confirming FUS-mediated opening of the BBB and the BTB (Figure 1.D; Table S2). No significant difference in Gd-DOTA enhancement was observed between the two FUS groups (FUS + carboplatin vs. FUS + carboplatin + RT) or between the non-FUS groups in any region (Figure S3; Table S3).

To assess acute safety, animals underwent MRI 48 h after the treatment and were euthanized for histological analysis. T₂-weighted, ADC and DCE-MRI images revealed no signal abnormality in any group; peritumoral hyperintensity on T₂-weighted images and elevated ADC values around the tumor core reflect tumor infiltration and edema, and were observed across all groups, confirming that these changes were not attributable to treatment toxicity. H&E-stained brain sections confirmed the absence of microhemorrhages or significant tissue damage (Figure S4).

### FUS-mediated BBB/BTB opening enhances carboplatin delivery to the tumor and peritumoral brain

To quantify carboplatin accumulation following BBB/BTB opening, brains and peripheral organs were collected from 22 animals at 1 h or 24 h after i.v. carboplatin administration and analyzed by ICP-MS (N = 14) or SR-XRF (N = 8) (Figure 2.A).

**Figure 2.**
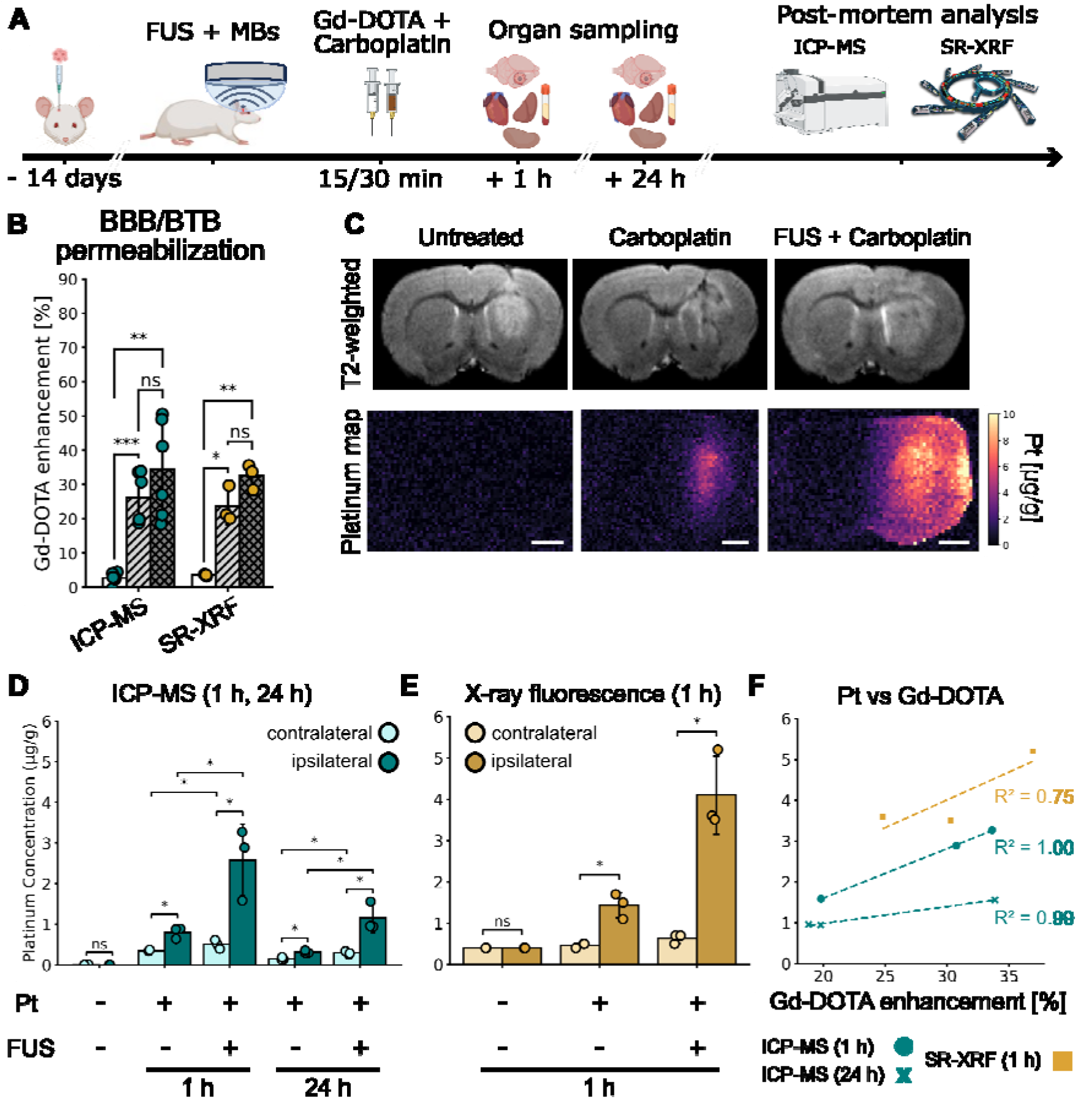
Platinum biodistribution following FUS-induced BBB/BTB opening. A) Experimental timeline for the biodistribution study. B) Gd-DOTA signal enhancement measured in the contralateral striatum (white), tumor periphery (hatched light gray), and tumor core (hatched dark gray) in groups analyzed by ICP-MS (green) and SR-XRF (yellow). *p*-values were calculated using two-sided paired t-tests (\**p* < 0.05, \*\**p* < 0.01, \*\*\**p* < 0.001; ns: not significant). C) Representative T2-weighted MRI and corresponding SR-XRF map for untreated, carboplatin-only, and FUS + carboplatin groups (scale bar, 2 mm). D) Platinum concentrations in the contralateral and tumor-bearing hemispheres measured by ICP-MS in untreated, carboplatin i.v., and FUS + carboplatin i.v. groups at 1 h and 24 h post-injection. *p*-values were calculated using two-sided paired t-tests (\**p* < 0.05; ns: not significant). E) Platinum concentrations measured by SR-XRF 1 h post-injection in untreated, carboplatin i.v., and FUS + carboplatin i.v. groups. *p*-values were derived from two-sided Mann–Whitney U-test (\**p* < 0.05; ns: not significant). F) Correlation between Gd-DOTA enhancement measured by MRI and platinum concentration in the tumor measured by ICP-MS (1 h (•) and 24 h (x), green) and SR-XRF (1 h, yellow square). Data are presented as mean ± SD.

Prior to comparing platinum concentrations between groups, we verified that Gd-DOTA enhancement was comparable between animals assigned to ICP-MS and SR-XRF cohorts, to rule out any experimental bias in BBB/BTB opening between analytical subgroups. Enhancement in the tumor core (34.6 ± 12.9% for ICP-MS; 32.6 ± 3.0% for SR-XRF) and in the peritumoral zone (26.1 ± 6.7% for ICP-MS; 23.7 ± 4.2% for SR-XRF) did not differ significantly between the two cohorts, confirming comparable BBB/BTB opening across animals assigned to each technique and to survival groups (Figure 2.B).

SR-XRF maps revealed different spatial distributions of platinum depending on FUS treatment. In carboplatin-only animals, platinum accumulation was essentially confined to the tumor core, consistent with the restricted transvascular transport across an intact BBB in the surrounding brain parenchyma (Figure 2.C; Figure S5.A). In contrast, FUS-treated animals displayed a broader distribution, extending well beyond the tumor core into the peritumoral brain zone (corresponding to the region permeabilized). Platinum concentrations in the contralateral hemisphere remain low (0.46 ± 0.06 µg/g and 0.63 ± 0.12 µg/g at 1h) and did not differ significantly between FUS-treated and non-FUS animals.

As measured by ICP-MS, FUS increased platinum concentration in the tumor-bearing hemisphere by 2.75-fold at 1 h and 3.28-fold at 24 h compared with non-FUS controls (Figure 2.D; Table S4). The ipsi-to-contralateral platinum ratio was 5.6 with FUS vs. 2.9 without FUS at 1 h and remained elevated at 24 h (4.3 vs. 2.5), indicating slow clearance from brain tissue. A modest 1.5- to 1.9-fold elevation in contralateral platinum was also observed in FUS-treated animals at 1 h and 24 h respectively, suggesting limited diffusion beyond the sonicated volume.

SR-XRF yielded comparable platinum concentrations (4.10 ± 0.95 µg/g with FUS vs. 1.43 ± 0.30 µg/g without FUS in the tumor-bearing hemisphere; Figure 2.E), supporting consistency of the two analytical approaches.

A linear correlation was observed between Gd-DOTA signal enhancement on T₁-weighted MRI and platinum concentration measured by ICP-MS at 1 h (R² = 1.00) and 24 h (R² = 0.99), and by SR-XRF at 1 h (Figure 2.F). This relationship was further supported at the spatial level, with the distribution of platinum measured by SR-XRF closely mirroring the pattern of Gd-DOTA enhancement on T₁-weighted MRI (Figure S5.B). Together, these findings indicate that contrast-enhanced MRI may provide a non-invasive marker of local carboplatin accumulation, with both spatial and quantitative correspondence.

Platinum concentrations in peripheral organs are detailed in Figure S6 and Table S4. The kidney showed the highest systemic accumulation at both timepoints, with progressive clearance between 1 h and 24 h. Small differences were measured at 1 and 24 h in kidney, heart and liver between FUS- and non-FUS-treated animals, yet no significant differences were measured.

### FUS-mediated carboplatin delivery combined with radiotherapy improves tumor control and prolongs survival

A total of six treatment groups were evaluated: untreated controls, radiotherapy alone (RT), carboplatin alone, carboplatin + RT, FUS + carboplatin, and FUS + carboplatin + RT. Baseline tumor volumes were comparable across all groups at treatment initiation with no significant difference (*p* = 0.89, one-way ANOVA). Kaplan–Meier analysis (Figure 3.A) showed that FUS + carboplatin + RT increased median survival by 33.0 days compared to untreated animals (60.0 vs. 27.0 days; 122% increase; *p* = 0.005, log-rank test). Median survival of the combined treatment group was also significantly longer than with FUS + carboplatin (60.0 vs. 28.5 days; 110% increase; *p* = 0.006), RT alone (60.0 vs. 41.5 days; 45% increase; *p* = 0.005) and carboplatin + RT (60.0 vs. 28.0 days; 114% increase; *p* = 0.049). In contrast, carboplatin + RT, carboplatin alone, and FUS + carboplatin alone did not significantly improve survival compared with untreated controls (median 28.0, 25.0, and 28.5 days; change by 4%, −7%, and 6%, respectively). Survival outcomes for all groups are summarized in Table 2. Complete pairwise log-rank test statistics are provided in Table S5.

**Figure 3.**
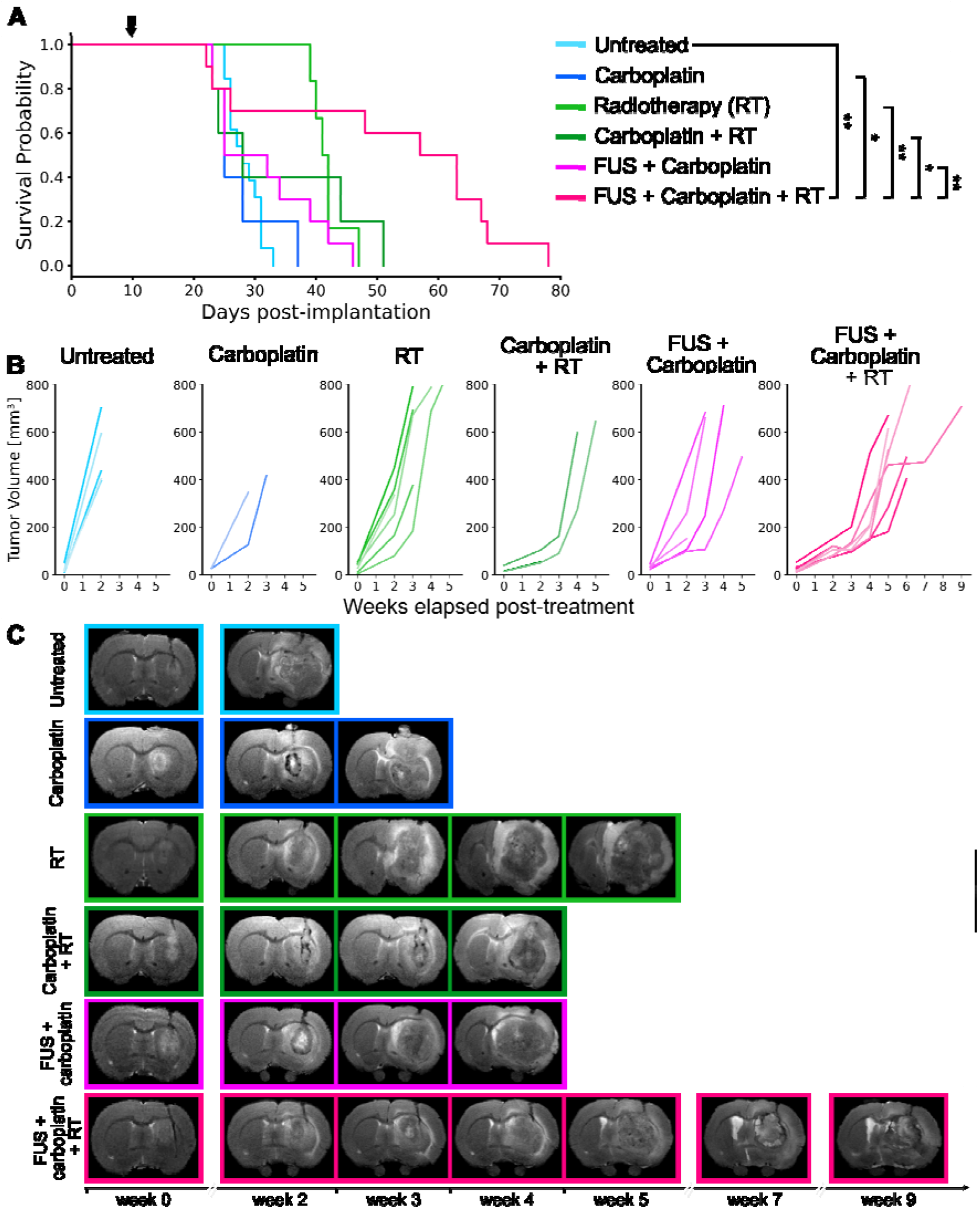
Survival and tumor growth. A) Kaplan–Meier survival curves for all treatment groups. The arrow indicates treatment initiation. P-values were derived from the log-rank test (*p* < 0.05: *; *p* < 0.01: **). B) Individual tumor volume evolution derived from T_2_-weighted hyperintense regions, shown for all animals in each treatment group throughout follow-up. C) T_2_-weighted anatomical MRIs acquired before treatment (day 0) and during follow-up, illustrating tumor growth in one representative animal per treatment group.

**Table 2.** Post-treatment survival outcomes in F98 glioma-bearing rats. Survival time was measured from the day of treatment. The increase in survival was calculated relative to the untreated group. N is the number of animals per group.

| **Group** | **N** | **Survival time [day]** |  |  | **Increased life span [%]** |  |
|---|---|---|---|---|---|---|
|  |  | Range | Mean ± SD | Median | Mean | Median |
| **Untreated** | 11 | 25 – 33 | 28.0 ± 2.8 | 27.0 | - | - |
| **Radiotherapy (RT)** | 6 | 39 – 47 | 41.8 ± 2.8 | 41.5 | 49 | 54 |
| **Carboplatin + RT** | 5 | 23 – 51 | 34.0 ± 12.7 | 28.0 | 21 | 4 |
| **Carboplatin** | 5 | 23 – 37 | 27.6 ± 5.5 | 25.0 | -1 | -7 |
| **FUS + carboplatin** | 10 | 23 - 46 | 31.4 ± 8.5 | 28.5 | 12 | 6 |
| **FUS + carboplatin + RT** | 10 | 22 – 78 | 51.5 ± 20.7 | 60.0 | 84 | 122 |

**Table 3.** Summary of MRI-derived and histological features of the identified tumor subregions.

| **Cluster** | **MRI-derived signature** | **Histological features** |
|---|---|---|
| **C1 (orange)** | Highest CBF values with increased AUC and prolonged TTP, consistent with increased vascular activity | Dense and irregular vascular networks with strong CD31 and ColIV staining, consistent with altered architecture; low Ki67 labeling |
| **C2**<br>**(cyan blue)** | High AUC values with prolonged TTP and reduced CBF, consistent with increased contrast accumulation, delayed contrast kinetics, and reduced perfusion | Partially disorganized vasculature with heterogeneous Ki67 labeling, consistent with vascular dysfunction and persistent cell proliferation |
| **C3 (yellow)** | MRI values closest to contralateral tissue, with moderately reduced CBF | Preserved tissue architecture and organized vasculature, consistent with relatively preserved peritumoral tissue or limited tumor infiltration |
| **C4 (pink)** | Low CBF with prolonged TTP and moderate permeability, reflecting severely impaired perfusion | Highly disorganized vasculature with fragmented vessel walls, sparse CD31 and Ki67 labeling, consistent with a poorly perfused and low-proliferative compartment |
| **C5 (purple)** | Highest ADC values, markedly reduced CBF, and increased permeability, consistent with tissue disruption and necrotic regions | Extensive necrosis, loss of tissue organization, abundant cellular debris, and minimal Ki67 labeling |

Individual changes in tumor volume derived from T₂-weighted MRI segmentation are shown in Figure 3.B, with representative images shown in Figure 3.C. Longitudinal changes in mean tumor volumes delineated on T2-weighted and T1-weghted images for all treatment groups are presented in Figure S7. Over the first two weeks post-treatment, untreated tumors expanded 19.0-fold in volume (506.3 ± 135.5 mm³ compared to 26.7 ± 17.9 mm³ at treatment initiation). Radiotherapy alone and carboplatin alone limited tumor growth to 8.5-fold (275.0 ± 135.3 mm³) and 8.2-fold (237.2 ± 155.3 mm³), respectively. Carboplatin + RT yielded the smallest increase in tumor volume at week 2 (3.0-fold; 70.5 ± 29.5 mm³), compared with 5.2-fold for FUS + carboplatin (155.1 ± 74.3 mm³) and 4.2-fold for FUS + carboplatin + RT (102.9 ± 20.1 mm³). At week 2, tumor volumes were significantly lower than in untreated controls in all treatment groups except carboplatin alone (all *p* < 0.05, Mann-Whitney U test). At week 3, mean tumor volumes were similar in the carboplatin + RT and FUS + carboplatin + RT groups (127.7 ± 51.6 and 124.9 ± 37.6 mm³, respectively; no significant difference). By week 4, mean tumor volume had increased to 436.2 ± 228.6 mm³ in the carboplatin + RT group, compared with 241.6 ± 131.2 mm³ in the FUS + carboplatin + RT group. Only FUS + carboplatin + RT group yielded significantly lower tumor volumes than RT alone at both weeks 3 and 4 (*p* = 0.017 and *p* = 0.022, respectively; Mann-Whitney U test) and tumor volumes in this group were also significantly lower than with FUS + carboplatin at week 3 (*p* = 0.042, Mann-Whitney U test). Similar growth was found on T1-weighted images following the injection of Gd-DOTA (Figure S7.B).

Among treated animals, early deaths, defined as deaths occurring before the upper bound of the survival range observed in untreated controls (<33 days), occurred exclusively in carboplatin-treated groups, including 2/5 in the carboplatin + RT, 5/10 in the FUS + carboplatin, and 3/10 in the FUS + carboplatin + RT. This pattern suggests a possible contribution of systemic carboplatin toxicity to early mortality, although the cause of death could not be established from these observations alone.

Body weight monitoring showed transient weight loss in all groups receiving carboplatin, while the radiotherapy-only and untreated groups maintained stable or increasing body weight throughout follow-up (Figure S8). Mean body weight reached its lowest value within the first two weeks in all carboplatin-treated groups: carboplatin alone at day 8 (−5.5%), FUS + carboplatin at day 4 (−6.4%), carboplatin + RT at day 12 (−7.7%), and FUS + carboplatin + RT at day 14 (−6.8%). Mean body weight returned to baseline in all these groups by days 19–20. No significant differences in weight loss were observed between FUS and non-FUS groups receiving the same chemotherapy regimen (FUS + carboplatin vs. carboplatin alone; FUS + carboplatin + RT vs. carboplatin + RT; all *p* > 0.05). These observations suggest that transient weight loss was primarily associated with systemic carboplatin exposure, although an additional effect of FUS cannot be excluded.

### Longitudinal multiparametric MRI revealed reduced expansion of high-ADC regions and transient increase in perfusion in the triple-combination group

Treatment response and tumor evolution were monitored through longitudinal quantification of ADC, CBF, and BBB permeability for up to 9 weeks post-treatment. Representative images from each treatment group are shown in Figure 4.A. These measurements revealed marked intratumoral heterogeneity and characterized changes in diffusion, perfusion, and permeability over time across brain regions and treatment groups.

**Figure 4.**
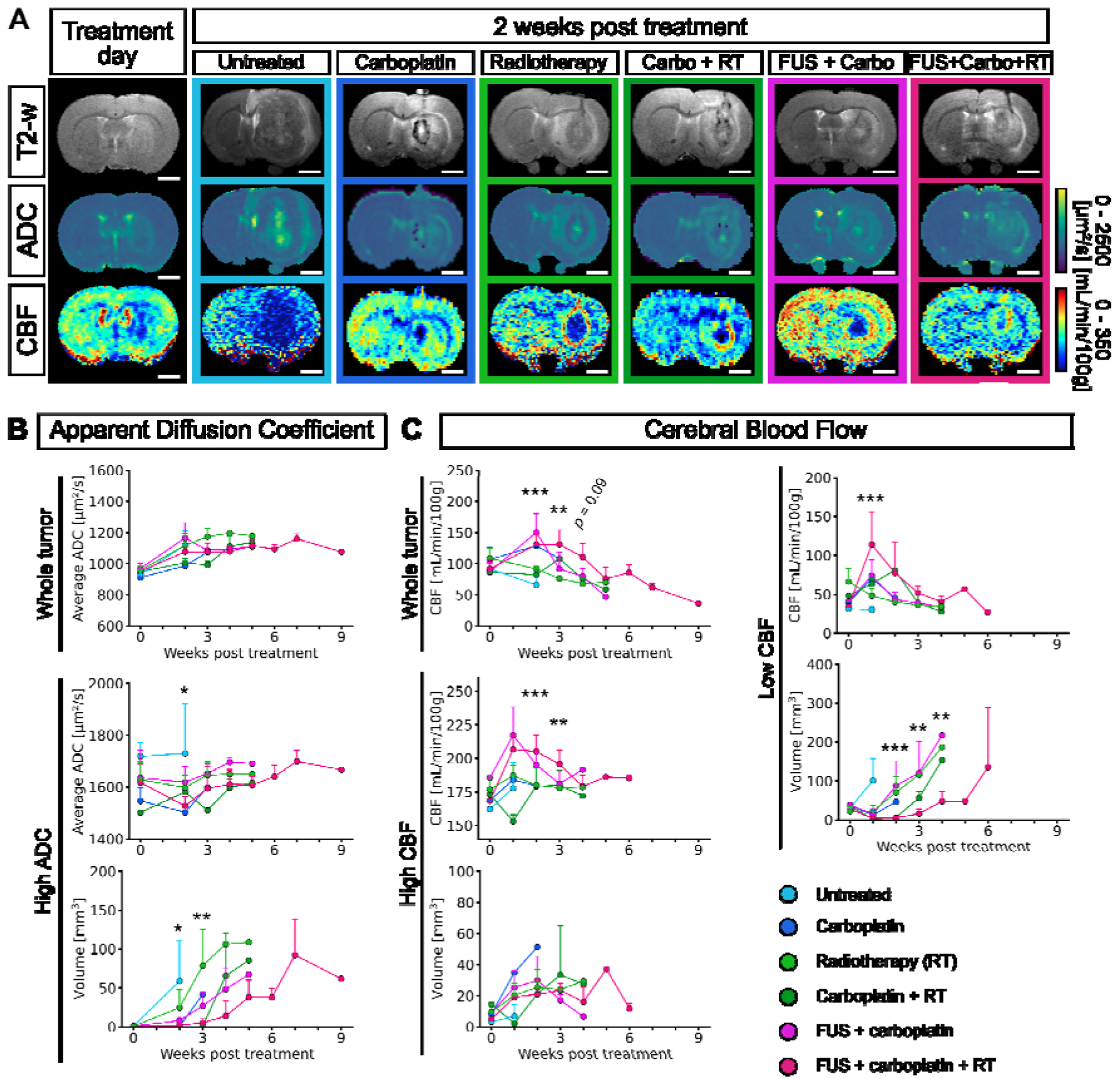
Longitudinal multiparametric MRI monitoring of tumor evolution across treatment groups. A) Representative T₂-weighted anatomical MR images, ADC maps, and CBF maps from each treatment group, acquired two weeks after treatment (scale bar, 2.5 mm). B) Longitudinal quantification of ADC values in the whole tumor volume (top), the high ADC zone (middle) and its volume evolution (bottom). C) Longitudinal CBF quantification in the whole tumor (top left panel), high CBF zone (bottom left panel), and low CBF zone (right panel). P-values were derived from one-way ANOVA (\**p* < 0.05, \*\**p* < 0.01, \*\*\**p* < 0.001; ns: not significant). Data are presented as mean ± SD.

ADC mapping revealed three distinct compartments: a central core with high ADC values (ADC > 1200 µm²/s), consistent with necrotic or severely disrupted tissue regions [49,50]; an intermediate peritumoral zone with moderate ADC values (800 < ADC < 1000 µm²/s), potentially reflecting viable but less densely packed tumor cells [50,51]; and a peripheral edematous region (ADC > 900 µm²/s), consistent with peritumoral fluid accumulation or tumor infiltration [52]. CBF maps revealed two compartments: a hypoperfused central core (CBF < 100 mL/min/100 g), surrounded by a hyperperfused ring (CBF > 120 mL/min/100 g).

#### Apparent diffusion coefficient

Longitudinal whole-tumor ADC analysis showed an initial increase during the first three weeks after treatment, followed by a plateau in all treatment groups except the carboplatin + RT group, in which ADC values continued to increase (Figure 4.B, top panel). In the high-ADC region, mean ADC values ranged from 1500 to 1700 µm²/s across treatment groups (Figure 4.B, middle panel), suggesting similar diffusion characteristics within these regions despite differences in treatment. ADC values in the contralateral striatum remained stable over time (Figure S9.A).

High-ADC region volumes differed significantly between treatment groups at weeks 2 and 3 after treatment (*p* = 0.021 and *p* = 0.002, respectively; one-way ANOVA), Figure 4.B, bottom panel. At week 2, carboplatin-treated groups had relatively small high-ADC region volumes (carboplatin: 0.37 mm³ (N = 1); FUS + carboplatin: 7.97 ± 5.19 mm³; carboplatin + RT: 0.03 ± 0.04 mm³; FUS + carboplatin + RT: 1.70 ± 2.20 mm³), whereas larger volumes were observed in the RT-only (24.67 ± 22.86 mm³) and untreated (58.91 ± 51.21 mm³) groups (Figure S10.A). At week 3, the FUS + carboplatin + RT group (5.10 ± 5.30 mm³) and the carboplatin + RT group (0.45 ± 0.63 mm³) had significantly smaller high-ADC region volumes than the RT-only group (78.50 ± 46.50 mm³; *p* = 0.006 and *p* = 0.036, respectively, Mann-Whitney U tests). The FUS + carboplatin + RT group also had significantly smaller high-ADC region volumes than the FUS + carboplatin group (27.00 ± 17.60 mm³; *p* = 0.029, Mann-Whitney U test), Figure S10.B.

By week 4, the carboplatin + RT group showed a marked increase in high-ADC region volume (65.6 ± 54.9 mm³), whereas the triple combination group maintained a smaller mean volume (13.9 ± 19.2 mm³) (Figure S10.C). Throughout the four-week follow-up period, the mean high-ADC region volume in the triple combination group remained below 14 mm³, representing less than 6% of the total tumor volume at week 4 (13.9 ± 19.2 mm³ vs. 241.6 ± 131.2 mm³).

#### Cerebral blood flow

Whole-tumor CBF measured using perfusion MRI is shown in Figure 4.C (top left panel), with longitudinal trajectories for all treatment groups presented in Figure S11. CBF differed significantly among treatment groups at week 2 after treatment (*p* < 0.001, one-way ANOVA). The FUS + carboplatin + RT group had significantly higher CBF values than the untreated controls (*p* = 0.008), RT-only (*p* = 0.004) and carboplatin + RT (*p* = 0.036) groups (post-hoc pairwise comparisons, Table S6). Significant differences among groups were also observed at week 3 (*p* = 0.002, one-way ANOVA), when the FUS + carboplatin + RT group continued to show significantly higher CBF values than the RT-only group (*p* = 0.003, post-hoc pairwise comparison). Results of post-hoc pairwise comparisons for whole-tumor CBF at all timepoints are provided in Table S6.

Within the high-CBF region, CBF values differed significantly among treatment groups at week 2 after treatment (*p* < 0.001, one-way ANOVA; Figure 4.C, bottom left panel). CBF was significantly higher in both FUS-treated groups, FUS + carboplatin (217 ± 21 mL/min/100 g) and FUS + carboplatin + RT (207 ± 12 mL/min/100 g), than in the RT-only group (187 ± 8 mL/min/100 g; *p* = 0.019 and *p* = 0.017, respectively, post-hoc pairwise comparisons, Table S7). In contrast, the carboplatin + RT group had the lowest CBF values of all groups (153 ± 5 mL/min/100 g), significantly lower than those in both the triple combination group (*p* = 0.036) and the RT-only group (*p* = 0.024, post-hoc pairwise comparisons, Table S7). These findings suggest that FUS-mediated BBB/BTB opening was associated with preservation of peritumoral perfusion when carboplatin was combined with radiotherapy. This difference persisted at week 3, with CBF remaining significantly higher in the triple combination group than in the RT-only group (*p* = 0.006, post-hoc pairwise comparison, Table S7).

The volume of the hypoperfused core differed significantly between treatment groups at two weeks after treatment (*p* < 0.001, one-way ANOVA; Figure 4.C, right panel). Both groups receiving carboplatin combined with radiotherapy, namely carboplatin + RT (0.3 ± 0.5 mm³) and FUS + carboplatin + RT (0.5 ± 0.3 mm³), showed an almost absent hypoperfused core at two weeks, and very small volumes at three weeks. In contrast, the radiotherapy-only group displayed an increase in hypoperfused core volume from 9.9 ± 4.8 mm³ at week 2 to 35.4 ± 17.7 mm³ at week 3, while the untreated group reached 58.5 ± 37.2 mm³ at week 2. By week 4, the hypoperfused core volume had markedly expanded in the carboplatin + RT group (35.8 ± 27.3 mm³), whereas it remained limited in the triple combination group (7.4 ± 14.6 mm³). This pattern was associated with more limited tumor growth between weeks 3 and 4 in the triple combination group than in the carboplatin + RT group. CBF measurements in the contralateral striatum are shown in Figure S9.B. Detailed statistical analyses of CBF measurements are provided in Tables S6-S8.

Taken together, these MRI findings indicate that FUS treatments were associated with transient increases in intratumoral and peritumoral perfusion at two and three weeks after treatment, an effect not observed in the carboplatin + RT group. In addition, the FUS + carboplatin + RT group showed limited expansion of high-ADC and hypoperfused regions throughout the five-week follow-up period, whereas these regions markedly expanded in the carboplatin + RT group at week 4. These findings were associated with more limited tumor growth in the triple combination group.

### Voxel-wise clustering of multiparametric MRI identifies spatially distinct tumor subregions associated with histological features

To investigate spatial and temporal heterogeneity within the tumor microenvironment, a voxel-wise unsupervised k-means clustering approach was applied to the tumor volume ROI using five quantitative MRI parameters: ADC, CBF, T_1_ relaxation time, area under the curve (AUC) and time-to-peak (TTP) (Figure 5.A).

**Figure 5.**
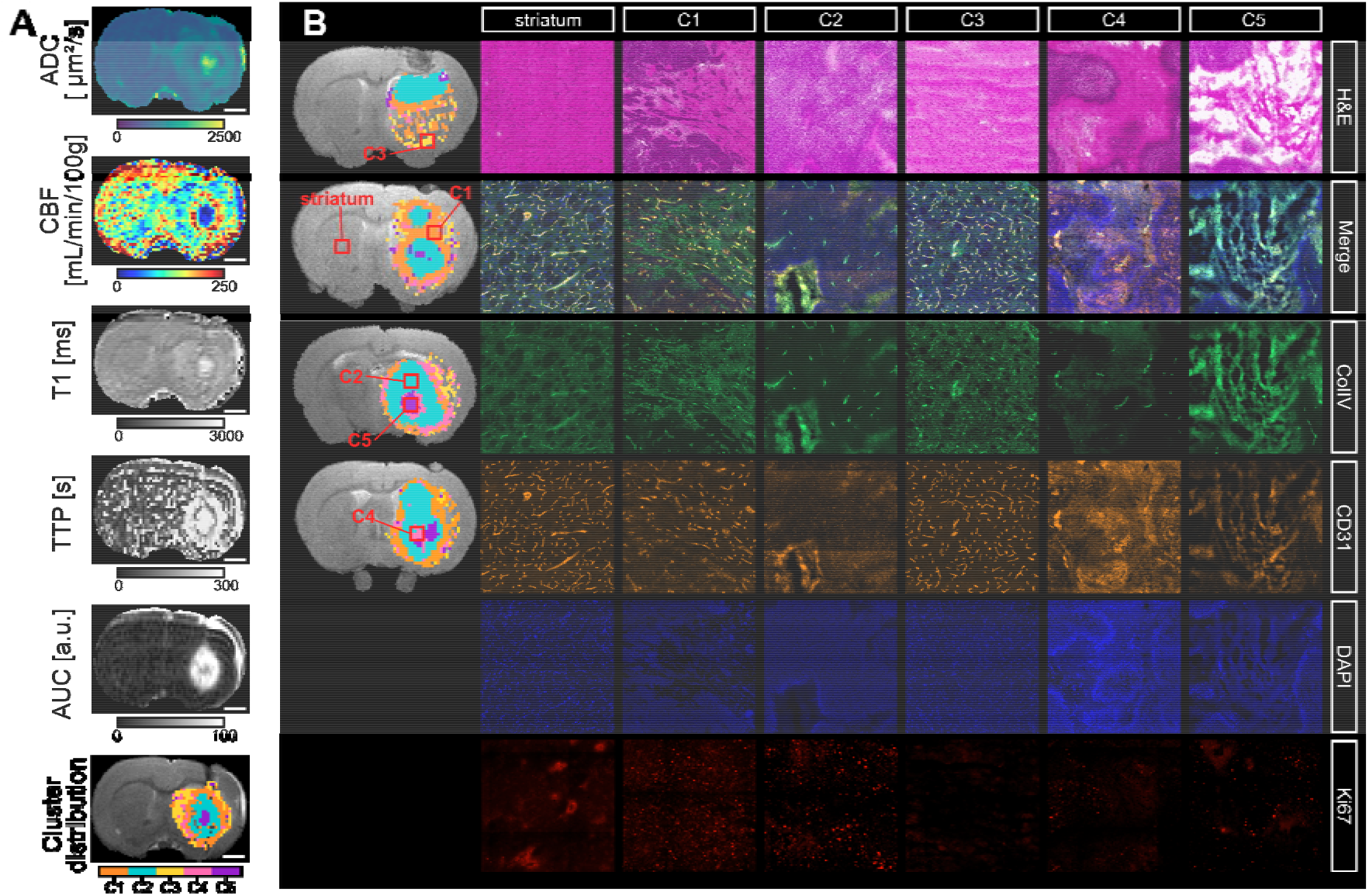
Multiparametric MRI-based clustering identifies tumor subregions with distinct histological features. A) Parametric MRI maps from a representative animal, including ADC, CBF, T_1_ relaxation time, and DCE-MRI-derived AUC and TTP. The spatial clustering map generated by voxel-wise k-means analysis of these parameters is overlaid on the anatomical T_2_-weighted image (scale bar, 2 mm). B) Histological and immunofluorescence images from selected brain regions corresponding to the contralateral striatum and representative cluster locations (C1 to C5). From top to bottom: H&E staining, merged fluorescence images (DAPI, CD31, ColIV), and individual channels showing CD31, ColIV, DAPI and Ki67 staining. Nuclei are labeled with DAPI (blue), blood vessels with CD31 (orange), vascular basement membranes with ColIV (green), and proliferating cells with Ki67 (red). All histological images in panel B represent a 1000 × 1000 µm^2^ field of view. Whole-slide images are presented in Figure S13.

Prior to clustering, all parametric maps were co-registered and standardized to ensure spatial alignment and comparability across MRI parameters. K-means clustering identified five distinct clusters within the tumor volume (C1 to C5). These accounted for 17.8%, 17.6%, 21.5%, 27.6%, and 15.6% of all voxels included in the analysis, respectively (Figure S12.A). Euclidean distance analysis between cluster centroids demonstrated clear separation between the identified clusters, supporting robust tumor sub-region delineation (Figure S12.B). Visualization of all tumor voxels in a low-dimensional UMAP (Uniform Manifold Approximation and Projection) embedding further illustrated the separability of the five subregions in the multiparametric MRI feature space (Figure S12.C). Table S9 summarizes the mean values of MRI-derived parameters for each cluster and for the contralateral region, which was used as a reference.

To support the biological interpretation of the MRI-based clustering, H&E staining and immunofluorescence analyses were performed on selected brain regions corresponding to representative clusters (Figure 5.B). H&E staining revealed histological features consistent with the tissue characteristics identified by MRI-derived maps.

Cluster 1 (C1, orange) displayed markedly elevated CBF values compared to the contralateral region (158.7 ± 59.8 vs. 130.8 ± 37.8 mL/min/100 g), increased AUC, and prolonged TTP, consistent with regions of high vascular activity and altered contrast kinetics. C1 was predominantly detected at the tumor periphery and was particularly prominent in FUS-treated animals, suggesting an association between FUS exposure and enhanced vascular remodeling rather than a simple physiological fluctuation. Histologically, C1 regions contained dense and irregular vascular networks, supported by strong CD31 and ColIV immunoreactivity, consistent with altered vascular architecture and increased vascular density. These regions exhibited low Ki67 labeling, indicating limited cell proliferation despite increased vascular activity.

Cluster 2 (C2, cyan blue) was primarily distinguished by high AUC values together with prolonged TTP values, consistent with increased contrast accumulation and delayed contrast kinetics, while CBF was lower than in the contralateral striatum. This profile suggests the presence of vessels with altered permeability and reduced perfusion. C2 regions were primarily detected within the tumor core during early stages and subsequently extended toward peripheral areas between clusters 3 and 5 as the tumor progressed. Histologically, these areas showed partially disorganized vascular structures and heterogeneous Ki67 labeling, supporting the MRI-based interpretation of vascular dysfunction associated with low but persistent cell proliferation.

Cluster 3 (C3, yellow) was characterized by MRI values closest to those of the contralateral brain, although with moderately reduced perfusion. It was predominantly located at the tumor periphery, suggesting regions with limited alterations in tissue microstructure or perfusion properties. Histologically, C3 regions exhibited preserved tissue architecture and organized vasculature, consistent with relatively preserved peritumoral tissue or areas with limited tumor infiltration, frequently located at the border of the T2-weighted hyperintense region. These findings highlight the limitations of conventional MRI-based segmentation methods for accurately delineating tumor tissue boundaries.

Cluster 4 (C4, pink) was characterized by low CBF, prolonged TTP, and moderate contrast accumulation, representing poorly perfused regions with dysfunctional vasculature. These areas likely correspond to vascular territories with severely impaired blood flow and limited cell proliferation. Gadolinium accumulation in these regions may have been flow-limited, potentially leading to an underestimation of vascular permeability despite extensive BBB/BTB disruption. Consistent with the MRI profile, C4 regions exhibited highly disorganized vascular structures, with fragmented vessel walls, as revealed by ColIV staining, together with sparse CD31 and Ki67 labeling. These histological features support the interpretation of C4 as a severely hypoperfused, poorly proliferative tumor compartment.

Cluster 5 (C5, purple) was characterized by an 87% increase in ADC values compared with contralateral tissue (1403 ± 257 vs. 750 ± 61 µm²/s), a 66% reduction in CBF, and increased contrast accumulation, reflecting impaired perfusion and tissue disruption, consistent with necrotic regions. C5 was predominantly localized within the tumor core, particularly at advanced disease stages, and its prevalence increased over time as the tumor progressed, especially in untreated and RT-only groups (Figure 5.A). Increased vascular permeability in these regions likely reflected altered BBB/BTB integrity and passive accumulation of Gd-DOTA. Corresponding histological sections revealed extensive necrosis, loss of tissue organization, and minimal Ki67 labeling, consistent with reduced proliferative activity. Dense DAPI staining in these areas indicated abundant nuclear material and cellular debris associated with necrotic tissue and elevated ADC values.

Overall, histological analyses revealed tissue features consistent with those inferred from the MRI-derived clusters, supporting the biological relevance of the multiparametric MRI-based classification. These findings suggest that voxel-wise clustering of quantitative MRI parameters can resolve spatially distinct tumor habitats characterized by specific vascular, structural, and proliferative features.

Subsequently, we analyzed the temporal evolution of cluster composition across treatment groups (Figure 6.A; individual cluster trajectories per group are shown in Figure S12.D). In tumors treated with radiotherapy alone, cluster proportions remained relatively stable throughout the follow-up period. However, at the end of the observation period, a marked increase in the proportion of C5 was observed compared with the other treated groups. In contrast, untreated tumors exhibited a progressive increase in C4 and a concomitant decrease in C1, suggesting a transition towards a phenotype characterized by poor perfusion and vascular dysfunction. Conversely, animals treated with FUS + carboplatin or FUS + carboplatin + RT exhibited similar cluster distributions two weeks after treatment, characterized by a marked reduction in the proportion of C2 and C4 together with an increase in the proportion of C1. The most pronounced changes were observed in the triple combination group, which displayed changes in tumor microenvironment composition over time, as illustrated by the longitudinal cluster maps. Figure 6.B provides representative examples of the spatial distribution of clusters over time for each treatment group. A marked predominance of C1 within the tumor was observed in the FUS + carboplatin + RT group compared with all other groups at weeks 2 and 3 after treatment. Near the endpoint, a clear divergence emerged between the triple-combination group and the other treatment groups, which displayed similar spatial patterns characterized by a high proportion of C3 and C5 clusters within the tumor. Importantly, these spatially resolved changes in cluster composition were not apparent when considering whole-tumor ADC values alone, highlighting the added value of multiparametric MRI-based clustering for capturing treatment-induced changes in the tumor microenvironment.

**Figure 6.**
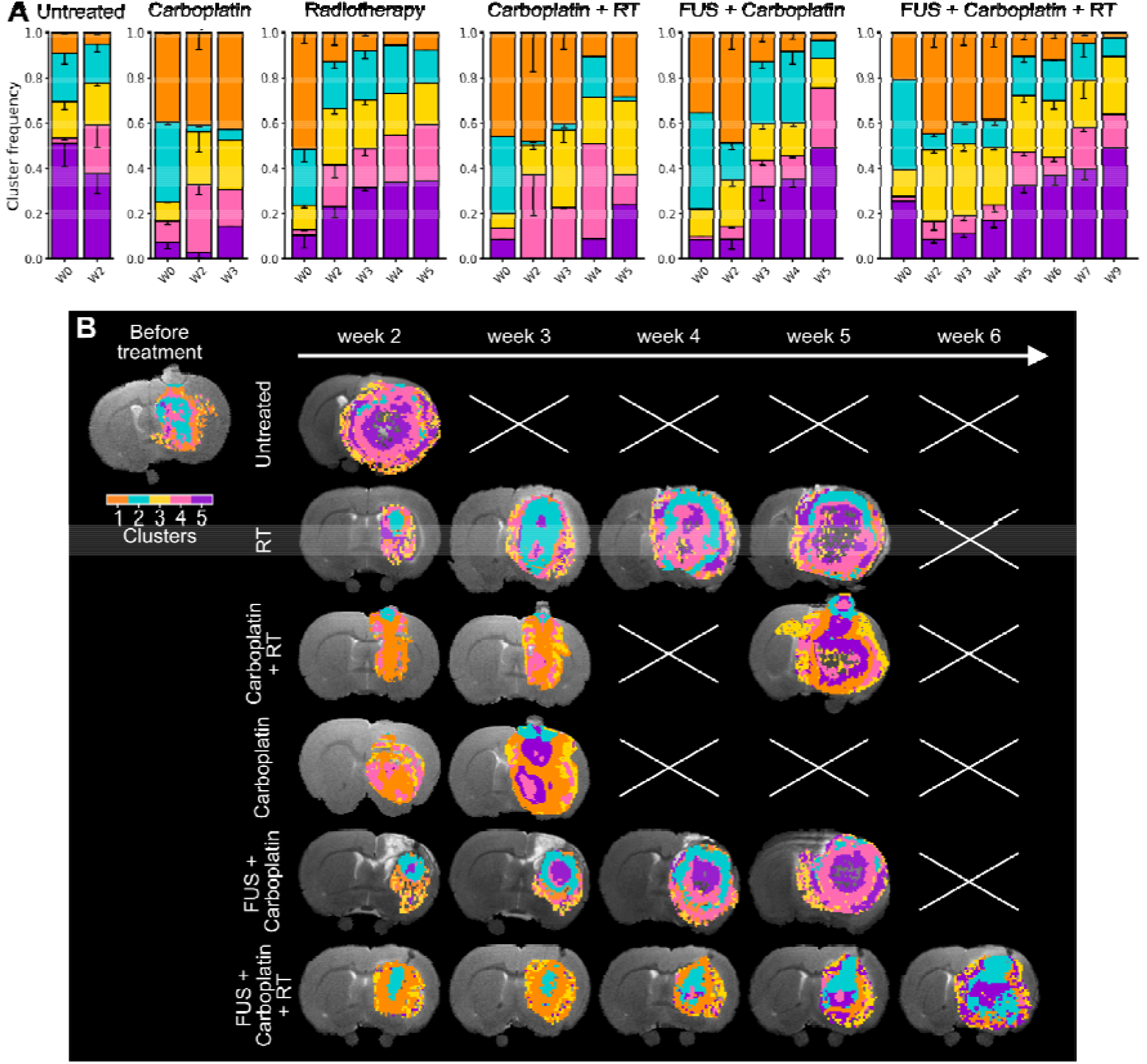
Longitudinal evolution of MRI-based clustering across treatment groups reveals treatment-associated changes in the microenvironmental composition. A) Relative frequencies of clusters C1 to C5 at different timepoints across treatment groups. Bars represent the proportion of tumor voxels assigned to each cluster at each imaging time point. B) Representative T_2_-weighted anatomical MRI images overlaid with cluster maps from individual animals in each treatment group at multiple follow-up timepoints. The color code corresponds to the five clusters derived from k-means analysis (C1–C5). Crossed boxes indicate missing data due to animal loss.

## Discussion

This study demonstrates that combining FUS-mediated carboplatin delivery with radiotherapy produces a significant therapeutic benefit in a syngeneic glioblastoma model, resulting in prolonged survival, reduced tumor growth, enhanced platinum delivery, and treatment-associated changes in the tumor microenvironment. Previous studies have demonstrated that FUS-mediated carboplatin delivery improves tumor control and prolongs survival in several preclinical glioma models [15,40,53]. These encouraging preclinical results have supported the translation of this approach into phase I/II clinical trials using implantable ultrasound devices, which have demonstrated its feasibility and safety [14,18]. In contrast, combining FUS with radiotherapy alone or with other chemotherapeutic agents, such as temozolomide or irinotecan, has not consistently improved survival in preclinical glioma models [24,25,54–56]. Our findings support the therapeutic potential of combining FUS-mediated carboplatin delivery with radiotherapy in this model.

In the F98 glioma model, platinum-based compounds, including carboplatin and cisplatin, have shown substantial anti-tumor activity, particularly when high intratumoral concentrations are achieved through local delivery approaches such as convection-enhanced delivery. Using this invasive approach, intratumoral platinum concentrations exceeding 10 µg/g (measured 1 h after infusion of 20 µg of carboplatin in 10 µL over 30 min [57]) were associated with significant survival benefits [58–60]. These findings highlight the importance of achieving sufficient intratumoral platinum exposure to obtain therapeutic efficacy. However, the clinical translation of these approaches remains limited by their invasive nature. FUS-mediated BBB/BTB disruption therefore represents a non-invasive and spatially targeted alternative to enhance platinum delivery to brain tumors.

In the present study, carboplatin administration following FUS-induced BBB/BTB opening resulted in mean platinum concentrations of 2.84 ± 0.86 µg/g and 4.10 ± 0.95 µg/g in the tumor region, as measured by ICP-MS and SR-XRF, respectively, 1 h post treatment. The differences observed between ICP-MS and SR-XRF measurements may reflect methodological differences, including tissue sampling (approximately 250 mg tissue fragments versus tissue sections of 40 µm, respectively) and uncertainties associated with absolute SR-XRF quantification, such as detector geometry and matrix effects.

Importantly, platinum remained detectable 24 h after carboplatin injection, corresponding to the time interval selected for radiotherapy delivery in our protocol. Following FUS-mediated BBB/BTB opening, platinum concentrations exceeding 1 µg/g were still detected in the tumor region. Over this interval, platinum levels decreased by approximately two-fold in brain tissue (including both tumor and contralateral regions with or without sonication), whereas plasma concentrations decreased 44- to 60-fold. These findings suggest prolonged tissue retention and/or delayed clearance of platinum relative to plasma [38,61]. Similar sustained intratumoral drug retention has been reported for ultrasound-mediated delivery of poorly diffusible agents such as doxorubicin, despite short permeability windows [62], and is consistent with the prolonged brain retention of carboplatin observed clinically by Habashi et al. following ultrasound-mediated BBB opening [63]. Overall, these results indicate that FUS-mediated delivery may be particularly advantageous for poorly diffusible agents such as platinum compounds, while emphasizing the importance of optimizing treatment parameters to maximize drug exposure and therapeutic efficacy.

Despite effective platinum delivery to the tumor-bearing hemisphere, FUS + carboplatin alone did not significantly improve survival compared with untreated animals (mean survival: 31.4 ± 8.5 vs. 28.0 ± 2.8 days). Signs of toxicity were observed in approximately 50% of treated animals, which displayed earlier mortality compared with untreated controls. Nevertheless, the survival range of FUS + carboplatin–treated animals (23 – 46 days) extended beyond that of the untreated animals (25 – 33 days), suggesting that a subset of animals may have derived therapeutic benefit despite the absence of an overall statistically significant survival improvement. Future studies could evaluate whether a lower carboplatin dose reduces toxicity, while maintaining therapeutic efficacy.

These results differ from those reported by McDannold et al., who reported a significant survival benefit following FUS + carboplatin treatment compared with untreated animals (41.7 ± 8.5 vs. 25.2 ± 1.5 days, respectively) [40]. In their study, mean carboplatin concentrations reached 21.6 µg/g, measured by LC-MS/MS in 3-mm tumor biopsies collected 1 h after injection, corresponding to 11.4 µg/g of Pt, which is approximately fourfold higher than the concentration measured by ICP-MS and 2.8-fold higher than that measured by SR-XRF in the present study. These differences likely reflect variations in experimental design, including the carboplatin dosing schedule, tumor implantation procedures, and treatment timing relative to tumor progression. Notably, McDannold et al. administered two carboplatin cycles at days 8 and 21 after tumor implantation, whereas the present study used a single administration around day 14.

Together, these findings suggest that enhanced platinum delivery with the single-dose regimen used here may not be sufficient to achieve durable tumor control. Rather, maximal therapeutic benefit may require combining enhanced drug delivery with complementary cytotoxic mechanisms. This concept is further supported by the comparative analysis presented in Table S10, which summarizes previously reported therapeutic strategies in the F98 glioma model, including BBB-opening approaches, drug delivery methods, radiotherapy, and their associated survival outcomes.

Among the treatment combinations evaluated in the present study, FUS-mediated carboplatin delivery combined with radiotherapy produced the greatest therapeutic benefit, resulting in sustained tumor growth delay for up to four weeks after treatment. When compared with the therapeutic approaches summarized in Table S10, this non-invasive combination achieved survival outcomes among the highest reported in the F98 glioma model, with a 122% increase in median survival compared to untreated controls. This efficacy compares favorably with other BBB-opening methods such as osmotic disruption [64], vasoactive agents [65], or FUS alone [40,66]. Although invasive local delivery methods, such as CED or osmotic pumps, may achieve greater survival benefits in some settings [59,67], their clinical applicability remains limited.

To further characterize the tissue changes associated with these therapeutic benefits, longitudinal multiparametric MRI was employed to non-invasively monitor the evolution of the tumor microenvironment across treatment groups. ADC and CBF, two established biomarkers of tumor cellularity and perfusion [27,68], revealed distinct treatment-associated dynamics. The combination of FUS + carboplatin + RT resulted in a sustained limitation of high-ADC and hypoperfused regions, suggesting delayed expansion of necrotic-like tumor compartments. Concurrently, an increase in CBF at two weeks after treatment in FUS + carboplatin-containing groups, suggested increased tumor perfusion and possible changes in vascular function. This vascular response may contribute to a more favorable microenvironment for radiotherapy efficacy, potentially through improved oxygen delivery [27,69], although this mechanism remains to be confirmed. In parallel, the stabilization of ADC values and the limited expansion of the low-CBF regions in the triple-treatment group suggest reduced microstructural deterioration and delayed necrotic progression, consistent with the observed reduction in necrotic volume.

Voxel-wise k-means clustering further captured treatment-induced changes in spatial tumor heterogeneity by identifying five distinct MRI-based subregions, in agreement with previous observations in preclinical glioma models [28,70]. Histological analyses supported the biological interpretation of these clusters, revealing differences in vascular architecture and density (CD31+/ColIV+), proliferative activity (Ki67 labeling), and tissue organization, including necrotic regions. This integrative imaging framework highlights the potential of clustering-based analysis as a spatially resolved biomarker of treatment response following FUS-mediated BBB/BTB opening, particularly for characterizing vascular and microenvironmental remodeling [22,37]. These findings highlight the potential translational value of multiparametric MRI clustering approaches for monitoring treatment response and potentially informing the timing of repeated FUS-mediated treatments. Similar data-driven approaches have successfully stratified glioblastoma patients according to progression risk and survival using diffusion and perfusion parameters [29,71], further supporting their potential for guiding adaptive FUS-based therapeutic strategies. Together with the histological characterization of the MRI-derived clusters, these observations suggest that FUS-containing treatments were associated with a transient vascular remodeling phenotype characterized by increased perfusion and dense vascular region.

Despite its promising efficacy and safety, this approach requires further optimization and safety assessment for clinical translation. The single-fraction radiotherapy regimen used here, although suitable for mechanistic evaluation, differs from current glioblastoma treatment protocols, which rely on fractionated treatments. Similarly, the absence of repeated carboplatin administration in the present study may have limited cumulative drug exposure and represents an important parameter to optimize in future translational protocols. Future investigations should determine whether repeated FUS-mediated carboplatin delivery combined with fractionated radiotherapy regimens can further enhance therapeutic efficacy while maintaining acceptable tolerability.

Beyond its role in enhancing drug delivery, FUS has also emerged as a potential immunomodulatory strategy, capable of reshaping the tumor immune microenvironment and enhancing responses to immune checkpoint inhibitors [72,73]. Exploring the combination of FUS-enhanced platinum delivery with immunotherapy therefore represents an attractive future direction [74,75]. While the F98 glioma model is well suited for evaluating cytotoxic treatment strategies owing to its infiltrative growth pattern and radioresistant phenotype, its limited immunogenicity may restrict its application in future immunotherapy studies.

Finally, the integration of multiparametric MRI with voxel-wise clustering provides a valuable framework for spatially characterizing tumor evolution and monitoring treatment-associated changes. Validation of these imaging biomarkers in clinical settings could support adaptive treatment strategies and patient stratification. In the present study, MRI-based analysis revealed differences in tumor composition that were not apparent from whole-tumor ADC values alone, suggesting that MRI-based monitoring may complement tumor volume measurements in the evaluation of therapeutic response and refinement of treatment planning. Overall, these findings demonstrate that transient, image-guided BBB/BTB opening enables effective carboplatin delivery and that its combination with radiotherapy improves therapeutic outcomes in the F98 glioma model. The combination of FUS-mediated drug delivery, longitudinal multiparametric MRI and spatial tumor characterization provides a comprehensive framework for evaluating future adaptive therapeutic strategies.

## Abbreviations

ADC: apparent diffusion coefficient
ANOVA: analysis of variance
AUC: area under the curve
BBB: blood-brain barrier
BTB: blood-tumor barrier
CBF: cerebral blood flow
CBCT: cone-beam computed tomography
CED: convection-enhanced delivery
ColIV: collagen IV
DAPI: 4T,6-diamidino-2-phenylindole
DCE-MRI: dynamic contrast-enhanced MRI
DWI: diffusion-weighted imaging
FUS: focused ultrasound
H&E: hematoxylin and eosin
ICP-MS: inductively coupled plasma mass spectrometry
i.v.: intravenous
MRI: magnetic resonance imaging
pCASL: pseudo-continuous arterial spin labeling
ROI: region of interest
SARRP: Small Animal Radiation Research Platform
SD: standard deviation
SR-XRF: synchrotron radiation X-ray fluorescence
TTFs: tumor-treating fields
TTP: time-to-peak
UMAP: Uniform Manifold Approximation and Projection

## Acknowledgments

The authors thank Nora Collomb (IRMaGe, La Tronche, France) for assistance with cell culture and tumor implantation, Pierre Juillard (Unité de Reconstitution Centralisée des Chimiothérapie, CHU Grenoble Alpes, France) for providing carboplatin chemotherapy, as well as Dr Caroline Boudou and Dr Barbara Fayard from Novitom for performing SR-XRF data reconstruction. This work was performed on the IRMaGe platform, a member of the France Life Imaging network. The Grenoble IRMaGe facility was partly funded by the French program “Investissement d’Avenir” run by the ‘Agence Nationale pour la Recherche’; grant ‘Infrastructure d’avenir en Biologie Santé’ (ANR-11-INBS-0006). The SAXO equipment was partly funded by: the ITMO Cancer of Aviesan within the framework of the 2021-2030 Cancer Control Strategy, with funds administrated by INSERM; The Rotary Club “Jetons le cancer 2022, H. Elleaume”; France Life Imaging, grant “ANR-11-INBS-0006”; Grenoble Alpes University (UGA); GEFLUC (H. Elleaume, 2022), Labex PRIMES ANR-11-LABX-0063, and the CALYPSO project, “INSERM Cancer PCSI 20CP082-00, H. Elleaume”. Part of the FUS equipment was sponsored by the Fédération pour la Recherche sur le Cerveau (FRC, Rotary Club, E. L. Barbier). This research was partly funded by ANR FUSBRAIN (Grant ANR-21-CE18-0019), a technological maturation project funded by C-Valo (PIA FNV Grant ANR-17-SATE-0003) and the Isère, Drôme and Loire committees of the Ligue Contre le Cancer (APP Auvergne Rhône Alpes - Saône et Loire 2024, H. Elleaume).

SR and SK were financially supported by ITMO Cancer of Aviesan within the framework of the 2021-2030 Cancer Control Strategy, with funds administered by INSERM (INCA, PCSI, 2023, IODA-Master, 23CP088-00, J.F. Adam).

We acknowledge the European Synchrotron Radiation Facility (ESRF) for provision of synchrotron beamtime under proposal number MD-1425 and we would like to thank Marco Di Michiel (ESRF) for assistance and support in using beamline ID15A. Data from the ID15A session are available under DOI 10.15151/ESRF-ES-1901512511.

This work was supported by the Photonic Imaging Center of the Grenoble Institute of Neuroscience (Univ Grenoble Alpes – Inserm U1216) which is part of the ISdV core facility and holds IBiSA certification. The authors are also grateful to the animal facility team at the Grenoble Institute of Neurosciences for their assistance with animal care and housing.

## Author contributions

SR, SK, SD, MDA and HE conducted the preclinical studies, including MR-guided FUS procedures, MRI follow-up, and radiotherapy. SR, SK, SD, MDA and HE prepared post-mortem tissue samples. SR and MDA processed ex vivo samples and performed histological staining and image acquisition. BC performed ICP-MS analysis. SR, SK, JFA, ALB, JLR, and HE performed SR-XRF acquisitions at the ESRF with assistance from M. Di Michiel (ESRF). SR-XRF data were reconstructed by NOVITOM (B. Fayard and C. Boudou, Grenoble, France). SR carried out all image analyses. SR and HE compiled and curated all data, prepared the figures, and wrote the manuscript. VS, BL, and ELB supported MRI analysis and clustering methodology. TA and AD prepared microbubble formulations and provided vials. SR, SD, SK, VS, JFA, ALB, AL and HE conceived and designed the study. ED, VS, AD, BL, ELB, JFA, ALB, JLR, and HE secured funding and supervised the project. All authors revised the manuscript critically, approved the final version, and agreed to be accountable for all aspects of the work.

